# Autophagy is required for lipid homeostasis during dark-induced senescence in Arabidopsis

**DOI:** 10.1101/2020.08.10.245092

**Authors:** Jessica A. S. Barros, Sahar Magen, Taly Lapidot-Cohen, Leah Rosental, Yariv Brotman, Wagner L. Araújo, Tamar Avin-Wittenberg

## Abstract

Autophagy is an evolutionarily conserved mechanism that mediates the degradation of cytoplasmic components in eukaryotic cells. In plants, autophagy has been extensively associated with the recycling of proteins during carbon starvation conditions. Even though lipids constitute a significant energy reserve, our understanding of the function of autophagy in the management of cell lipid reserves and components remains fragmented. To further investigate the significance of autophagy in lipid metabolism, we performed an extensive lipidomic characterization of Arabidopsis (*Arabidopsis thaliana*) autophagy mutants (*atg*) submitted to dark-induced senescence conditions. Our results revealed an altered lipid profile in *atg* mutants, suggesting that autophagy affects the homeostasis of multiple lipid components under dark-induced senescence. The acute degradation of chloroplast lipids coupled with the differential accumulation of triacylglycerols (TAGs) and plastoglobuli indicates an alternative metabolic reprogramming towards lipid storage in *atg* mutants. The imbalance of lipid metabolism compromises the production of cytosolic lipid droplets and the regulation of peroxisomal lipid oxidation pathways in *atg* mutants.

**One-sentence summary:** Autophagy is required for the mobilization of membrane lipid components and lipid droplet dynamics during extended darkness in *Arabidopsis*.

## Introduction

The precise control of energy homeostasis is crucial for plant survival, particularly under biotic and abiotic stress conditions. Under low energy availability, plants require a strategy other than carbohydrate oxidation to fully sustain energy production. Therefore, proteins, lipids, and chlorophylls can be catabolized instead of sugars to provide alternative substrates for the tricarboxylic acid (TCA) cycle. This assures the continued operation of the mitochondrial electron transport chain (Ishizaki et al., 2005; Kunz et al., 2009; Araujo et al., 2011; Hortensteiner & Krautler, 2011).

Among the alternative substrates of respiration in plants, lipids comprise a significant carbon reserve that represents an important alternative source of energy. Although in comparison to carbohydrates, the usage of lipids is less efficient, as assessed by the respiratory quotient, their intracellular storage and utilization are critical for the maintenance of cellular energy homeostasis (Araujo et al., 2011; Yang et al., 2018). Plants store lipids as the neutral lipid species TAG (Yang et al., 2018). In leaf mesophyll cells, TAGs are deposited in specialized storage compartments, namely lipid droplets (LDs) in the cytosol or plastoglobuli (PG) inside chloroplasts (Van wick et al., 2017). During nutrient deprivation, cellular lipid reserves are hydrolyzed into fatty acids (FAs), which are further catabolized during β-oxidation. Accordingly, mutants with dysfunctional β-oxidation, triggered by the disruption of the PEROXISOMAL ABC TRANSPORTER 1 (PXA1), exhibit hypersensitivity to extended darkness (Kunz et al., 2009). Moreover, several transcripts of the β-oxidation pathway are up-regulated during developmental leaf senescence (Troncoso-Ponce et al., 2013).

Autophagy, one of the primary cellular catabolic process, is involved in the degradation and recycling of cytoplasmic constituents into primary metabolites (Avin-Wittenberg et al., 2018). The autophagy mechanism is governed by autophagy-related (*ATG*) genes that are highly conserved among eukaryotes (Marshall and Vierstra, 2018). Plants with genetic disruption of *ATG* genes (hereafter *atg* mutants) have been well characterized for sensitivity to several stresses, particularly nutrient starvation (Avin-Wittenberg, 2018). The importance of autophagy in plants has been mostly associated with the recycling of proteins to amino acids under starvation conditions (Izumi et al., 2013; Li et al., 2014, Barros et al., 2017: Hirota et al., 2018; Mcloughin et al., 2018, 2020). By contrast, in mammals and yeast, autophagy has been largely associated not only with proteins but also with lipid metabolism (Elander et al., 2018; Barros et al., 2020). Accordingly, it was shown that autophagy supplies FAs for LD biogenesis as well as mediates the selective degradation of LDs in both mammals and yeast (Singh et al., 2009; Rambolt et al., 2015; Ngyugen et al., 2017).

A growing body of evidence suggests a dual role of autophagy in the lipid metabolism of alga and plants. Thus, in Chlamydomonas (*Chlamydomonas reinhardtii*), it has been postulated that autophagy participates in both TAG biosynthesis and LD formation (Couso et al., 2017; Pugkaew et al., 2017). In addition, microscopic evidence pointed to the autophagy-mediated engulfment of LDs in the algae *Auxenochlorella protothecoides* (Zhao et al., 2014) and *Micrasterias denticulata* (Schwarz et al., 2017). Interestingly, recent data also suggested that autophagy may mediate LD formation during an early stage of starvation while likely operating in LD degradation at the later stages of nitrogen starvation in Chlamydomonas (Tran et al., 2019).

In vascular plants, it was demonstrated that autophagy contributes to TAG synthesis via the turnover of organellar membranes in Arabidopsis under normal growth conditions (Fan et al., 2019). On the other hand, it was suggested the autophagy operates in the degradation of LDs by a selective microautophagy mechanism under extended darkness (Fan et al., 2019). Notably, this study was based on the analysis of Aradiposis mutants overexpressing the LD marker OLE1-GFP and knockout for SUGAR DEPENDENT-1 (SDP1) lipase, which are genotypes with altered LD and TAG metabolism (Fan et al., 2013, 2017; Barros et al., 2020). In maize (*Zea mays*), *atg12-1* mutant presented a higher LD number, which could reflect active LD assembly or disruption of LD catabolism (Mcloughin et al., 2020). It remains contentious, however, whether autophagy mediates the biogenesis or degradation of LDs in vegetative tissues (Elander et al., 2018; Barros et al., 2020; Masclaux-Daubresse et al., 2020). Although recent studies have clearly revealed an interplay between autophagy and lipid metabolism (Mcloughin et al.,2018, 2020; Have et al., 2019; Fan et al., 2019), relatively little is known about the role of autophagy in the mobilization of lipid components under energy limitation conditions.

Here, we investigated the significance of autophagy in lipid metabolism during dark-induced senescence in Arabidopsis. By performing an extensive lipidomic analysis of three independent *atg* mutants, we observed an overall impact of autophagy deficiency on all lipid classes analyzed and the free fatty acid (FFA) profile. The differential pattern of phospholipid levels and TAG accumulation was further coordinated with changes in LD number and PG size. In addition, analysis of β-oxidation genes revealed an induction of lipid oxidation pathways in *atg* mutants.

Collectively, our results demonstrate that autophagy is an essential component in the management of lipid reserves, affecting membrane lipid mobilization, the balance of LD and PG formation, and the regulation of β-oxidation pathways under dark-induced senescence.

## Results

### Differential lipid response of *atg* mutants under dark-induced senescence

To examine the involvement of autophagy in lipid remodeling during dark-induced senescence, we analyzed three previously described Arabidopsis autophagy mutants: *atg5-1* and *atg7-2* with full inhibition of autophagy (Chung et al., 2010); and *atg9-4* mutant that displayed a milder reduction of the autophagic process (Shin et al., 2014). The experimental setup consisted of the submission of 4-week-old plants to extended darkness conditions, a system widely used to induce energetic stress in plants (Ishizaki et al., 2005; Araujo et al., 2010; Barros et al., 2017).

To further investigate the impact of autophagy disruption on lipid homeostasis during dark-induced senescence, we performed Liquid Chromatography–Mass Spectrometry lipidomic analysis of *atg5-1, atg7-2, atg9-4*, and their corresponding wild-type (WT) plants. We were able to identify 119 lipid species of the following classes: Phosphatidylcholine (PC), Phosphatidylethanolamine (PE), Diacylglycerol (DAG), Triacylglycerol (TAG), Monogalactosyldiacylglycerol (MGDG), and Digalactosyldiacylglycerol (DGDG). We first performed a principal component analysis (PCA) based on the lipid profiles of samples from control and 6 days (d) of darkness (Fig. 1). Most of the variance of the dataset was covered by the first two principal components, explaining 69.1% of the overall data variance (57.8% and 11.3% for PC1 and PC2, respectively). This fingerprint analysis allowed us to obtain a general picture, discerning the main differences between the darkness treatment and the *atg* mutant lines. Briefly, it revealed that under control conditions (before darkness treatment), all genotypes clustered together; however, a clear separation according to the treatment and genotype was observed following darkness, wherein *atg7-2* and *atg5-1* were disconnected of the WT samples (Fig. 1). Interestingly, *atg9-4* presented an intermediary distribution, sharing certain metabolic similarities with WT and the other *atg* genotypes. The analysis of all data points also highlighted the delayed response of *atg9-4*, which after 6d and 9d of darkness, was more closely related to *atg5-1* and *atg7-2* after 3d of darkness conditions (Supplemental Figure S1).

**Figure 1.**
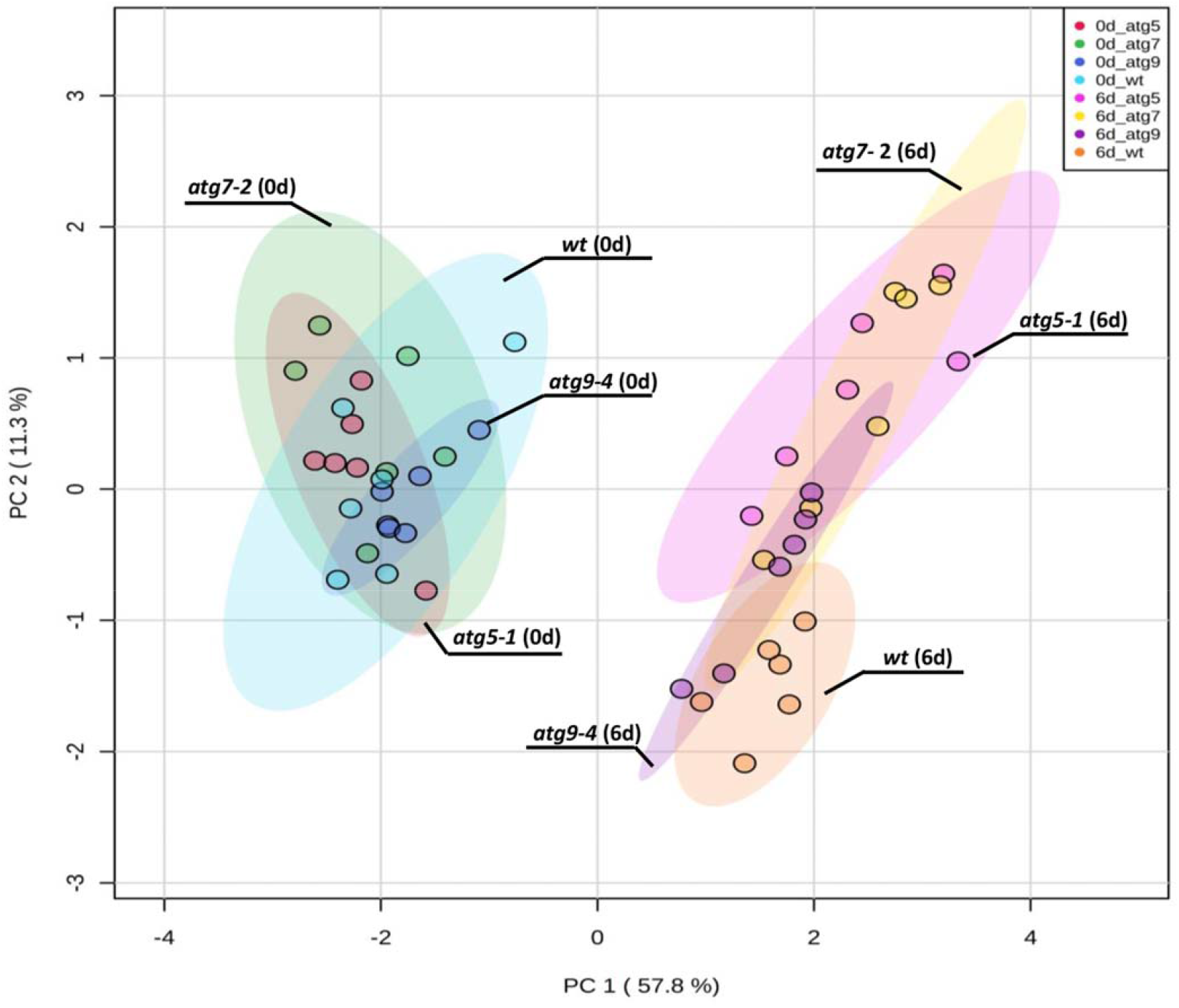
Differential lipid response of *atg* mutants under dark-induced senescence. Principal component analysis (PCA) was performed using the lipid data obtained in samples from 4-week-old Arabidopsis plants immediately before lights were turned off (0 d) and after further treatment for 6d in darkness. Three loss-of-function mutants of the autophagy pathway, namely *atg5-1, atg7-2, atg9-4* and their respective wild type (WT) were analyzed.

Our results are in close agreement with the milder impact of the *ATG9* disruption on metabolic features previously reported under dark-induced senescence (Barros et al., 2017). This response was related to residual autophagy activity observed in *atg9* lines (Yoshimoto et al., 2004; Shin et al., 2014), which explains the intermediary lipid response of *atg9-4* genotype. The information of the individual variables that contributed most to PC1 and PC2 (i.e., the lipids with the main impact on the variance of the dataset) is additionally available in Supplemental Table S1. Several lipids accounted for the main changes observed in the PCA. Briefly, PC1 was mainly comprised of MGDG 34:4, MGDG 34:3, PC 38:4(1) and MGDG 34:1 underlining the separation of darkened and control samples. On the other hand, TAG 54:9, MGDG 32:3 (1), and TAG 52:6 were the main lipids responsible for changes in PC2 (Supplemental Table 1), contributing to the differences between the genotypes under darkness (Fig. 1).

### The lack of autophagy affects the dynamics of phospholipids during dark-induced senescence

In order to obtain a more detailed characterization of the lipid composition of *atg* mutants following extended darkness, we analyzed each lipid class separately. We were able to identify the two dominant structural phospholipid species, namely PC and PE. These lipids are components of the plasma membrane, in which PC is associated with the outer leaflet, and PE is concentrated in the inner leaflet. PC and PE are also precursors of lipid mediators, such as phosphatidic acid (PA) and diacylglycerol (DAG) (Wang et al., 2006; Testerink and Munnik, 2011). Additionally, PE plays a crucial role in autophagosome formation (Soto-Burgo et al., 2018; Barros et al., 2020). Interestingly, a slight accumulation of both PC and PE was observed in *atg5-1* and *atg7-2* mutants before darkness treatment (control conditions, Fig. 2 – 0d). The difference of PC levels under control conditions was marked by the accumulation of 36C PC in *atg5-1* and *atg7-2* mutants, whereas no significant changes were observed in *atg9-4* at this time point (Fig. 2A – 0d). Extended darkness triggered a mixed response of different PC species. Thus, the levels of 32C and 34C mostly increased in WT plants under darkness. However, *atg5-1* and *atg7-2* presented lower accumulation of 34C PC species that contain a combination of 18C and 16C polyunsaturated acyl species, such as 34:5- (18:3-16:2-) and 34:6- (18:3-16:3) during dark treatment. The PC containing FAs with longer chains, 36C and 38C, were generally reduced in response to extended darkness. However, *atg5-1* and *atg7-2* presented a milder reduction of 36C PC species after 3d and 6d of darkness, whereas the 38C species (38:3(2), 38:4, 38:5, 38:6) were more reduced in these genotypes after 6d and 9d of darkness (Fig 2A).

**Figure 2.**
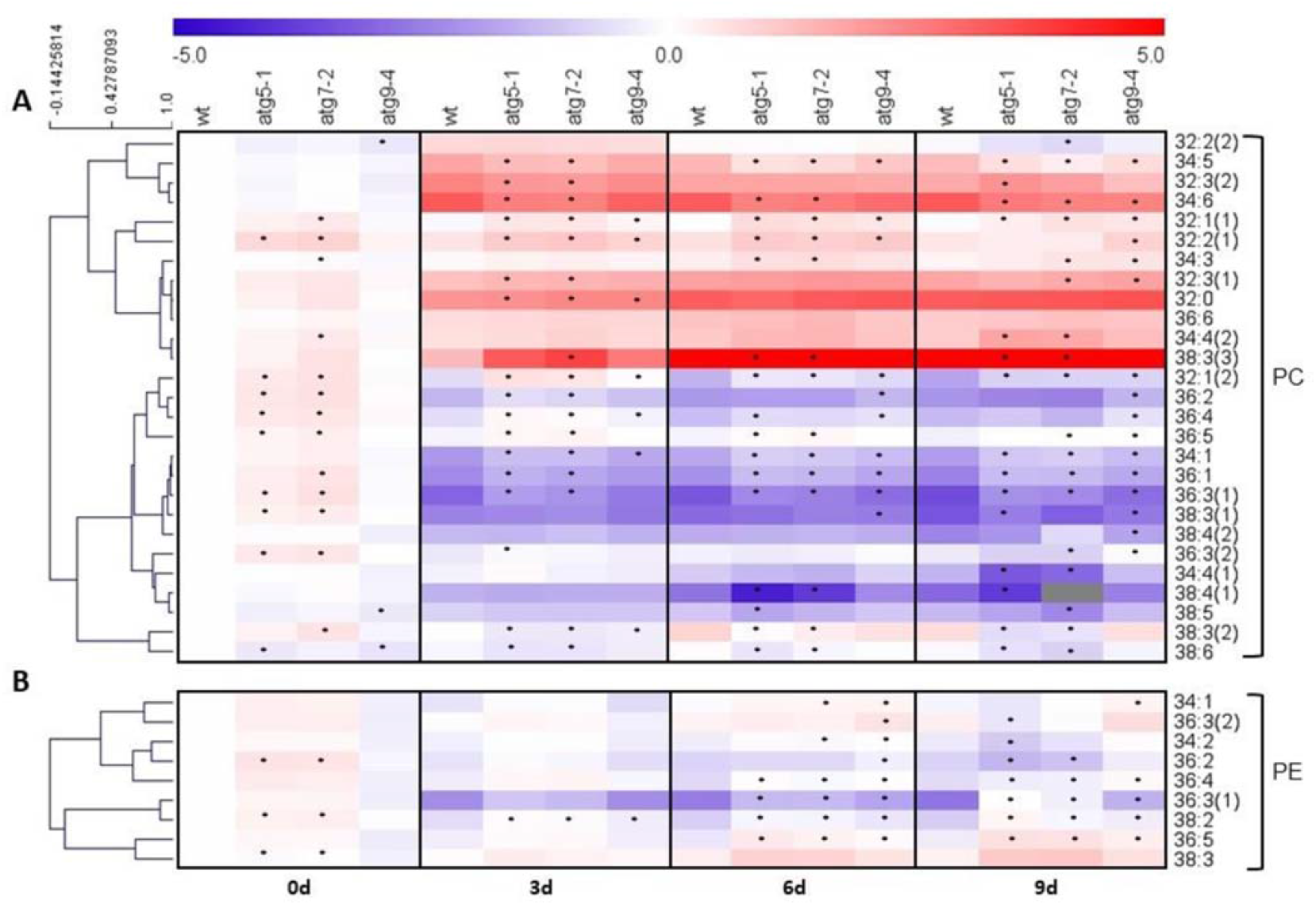
*atg* mutants present altered phospholipid levels under control and extended darkness conditions. Hierarchical clustering of (A) Phosphatidylcholine (PC) and (B) Phosphatidylethanolamine (PE). Values plotted are log_2_ fold change. Distances were measured by Pearson correlation using MeV 4.9.0 software. Data were normalized to the mean response calculated for the 0-d dark treated leaves of the wild type (WT). Values presented are means ± SE of five-six biological replicates per genotype; an asterisk (*) designates values that were determined by Student’s t-test to be significantly different (*P* < 0.05) from WT at each time point analyzed.

Following dark treatment, the majority of PE species decreased in WT plants; however, this reduction was milder in *atg* mutant lines after 6d of darkness (Fig. 2B). On the other hand, after 9d of darkness, species such as PE 36:2, 34:2, and 36:3(2) were highly reduced in *atg5-1* and *atg7-2* compared to WT. Surprisingly, the PE species 36:5 and 38:3 presented a contrasting pattern, accumulating after 6d and 9d of darkness in *atg* mutants (Fig. 2B).

### Impairment of autophagy accelerates the loss of chloroplast lipids

Since chloroplast lipids are subjected to constant turnover during senescence, we next evaluated the predominant lipid constituents of chloroplast membranes, MGDG and DGDG. An apparent reduction in more than 90% of the detected MGDGs and DGDGs was observed in *atg* mutants exposed to dark treatment (Fig. 3). This response was intensified throughout the dark treatment, and thus the levels of some species were below the detection threshold after 9d. Although *atg9-4* mutants followed the same pattern, the levels of MGDG and DGDG were less reduced in comparison to *atg5-1* and *atg7-2* mutants (Fig. 3). The overall reduction of chloroplast lipids reported here is consistent with the more severe phenotype described previously for *atg5-1* and *atg7-2* mutants lines under extended darkness (Barros et al., 2017). Interestingly, slightly lower levels of unsaturated MGDG and DGDG were also observed in *atg* mutants under control conditions, highlighting the potential role of autophagy in basal chloroplast maintenance, as recently verified in different studies using both Arabidopsis *atg5* and maize *atg12* mutants (McLoughlin et al., 2018, 2020; Have et al., 2019).

**Figure 3.**
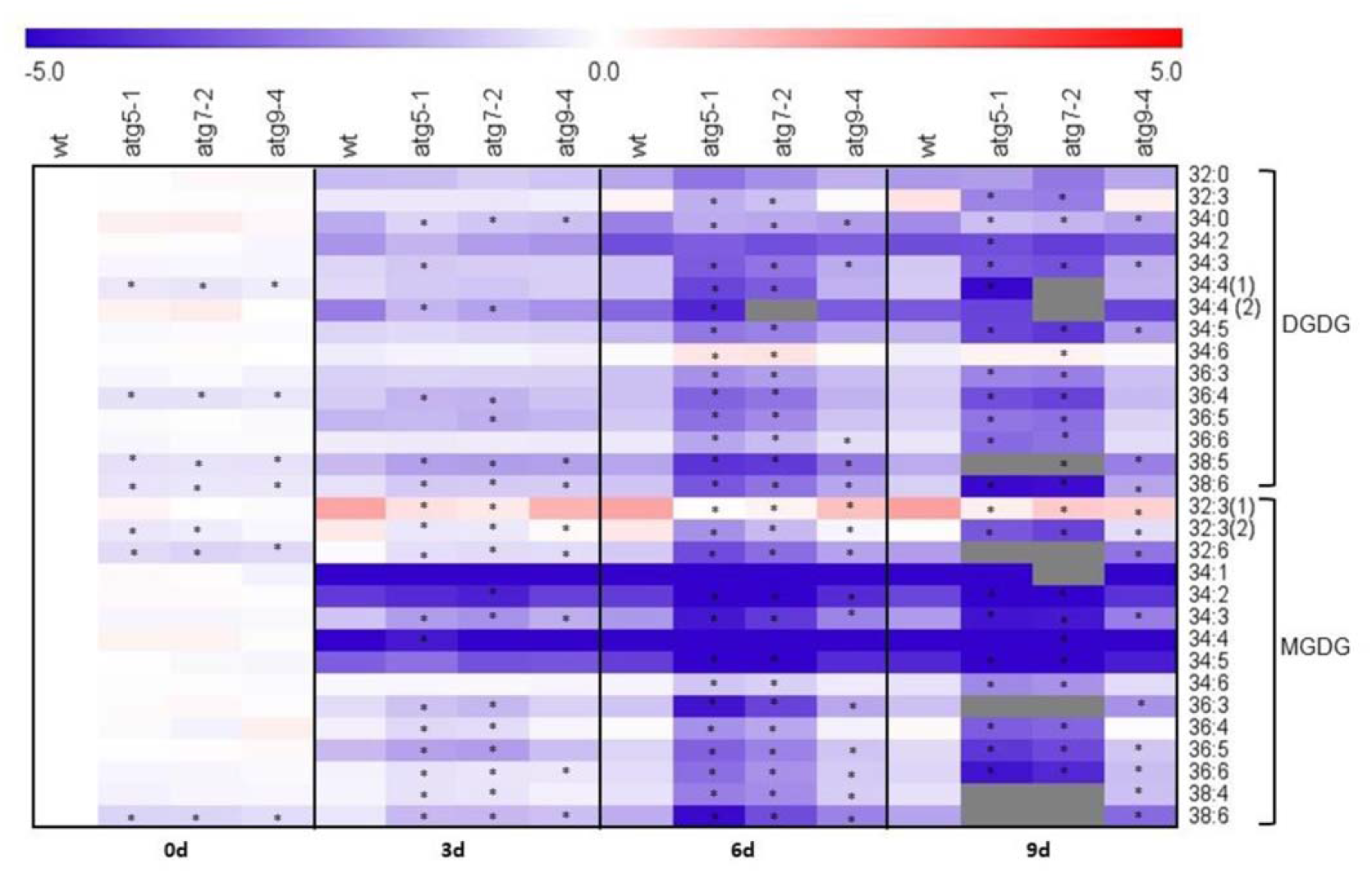
Chloroplast lipids are substantially reduced in *atg* mutants during dark-induced senescence. Monogalactosyldiacylglycerol (MGDG) and Digalactosyldiacylglycerol (DGDG) values plotted are log_2_ fold change. Data were normalized to the mean response calculated for the 0-d dark treated leaves of the wild type (WT). Values presented are means ± SE of five-six biological replicates per genotype; an asterisk (*) designate values that were determined by the Student’s t-test to be significantly different (*P* < 0.05) from WT at each time point analyzed.

### Disruption of autophagy compromises TAG and LD response under dark-induced senescence

The analysis of neutral lipids revealed a general increase in TAG levels during extended darkness. TAGs are composed of three FAs esterified to a glycerol backbone (Yang et al., 2018). The dominating accumulated TAG species during dark-induced senescence harbor three FA molecule with 18C with two or three double bonds (54:7 to 54:9 in Fig. 4A), or two FA molecules with 18C with two to three double bonds and 16:3 or 16:0 as a third FA (52:5 to 52:9 in Fig. 4A). Unsaturated C18 FAs (oleic acid, 18:1; linoleic acid, 18:2, and α-linolenic acid, 18:3), palmitic acid (16:0) and unsaturated C16 FAs (16:1 and 16:3) are highly abundant in chloroplasts of Arabidopsis (Hölzl and Dörmann, 2019). Our data indicate that the accumulated TAG species are most likely derived from plastidial lipids. Interestingly, the increase in chloroplast-derived TAG species was less pronounced in both *atg5-1* and *atg7-2* mutants after 3d and 6d of darkness. Additionally, an increase of other TAG species was observed from 6d of dark treatment onwards in *atg5-1* and *atg7-2* mutants, resulting in general TAG accumulation during the late period of darkness (Fig. 4A).

**Figure 4.**
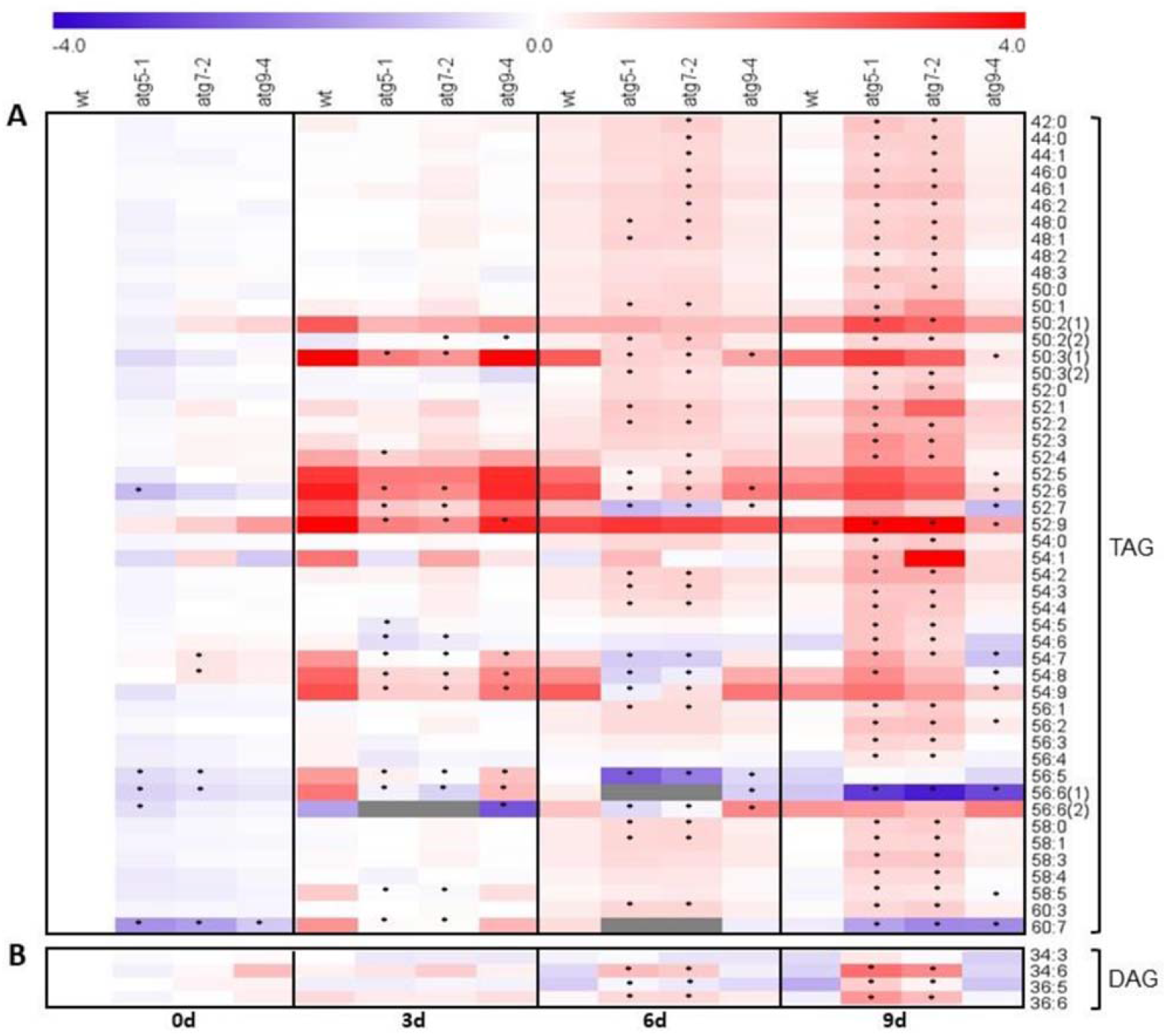
Distinct accumulation of neutral lipids in *atg* mutants during dark-induced senescence. (A) Triacylglycerol (TAG) and (B) Diacylglycerol (DAG). Values plotted are log_2_ fold change. Data were normalized to the mean response calculated for the 0-d dark treated leaves of the wild type (WT). Values presented are means ± SE of five-six biological replicates per genotype; an asterisk (*) designates values that were determined by Student’s t-test to be significantly different (*P* < 0.05) from WT each time point analyzed.

DAG is a central metabolite in plant lipid metabolism, serving as an intermediate of both TAG and galactolipids (Muthan et al., 2013). Similar to the observed for TAGs, the four DAG molecules identified (34:3, 34:6, 36:5, 36:6) were composed of unsaturated FAs, containing at least one unit of 16:2, 16:3 or 18:3 plastid-derived FA (Fig. 4B). Although no significant differences were observed during the early stages of dark treatment, DAGs started to accumulate 6d after the onset of dark treatment in *atg5-1* and *atg7-2* mutants. At the same time, WT and *atg9-4* plants showed a trend of reduction (Fig. 4B). The accumulation of unsaturated DAG species is likely a result of the degradation of chloroplast membranes in *atg* mutants serving as a precursor for the general TAG accumulation in the late stages of dark-induced senescence.

TAG is the central lipid reserve of the plant cell, stored in LDs (Pyc et al., 2017). To further investigate how the altered levels of TAGs in *atg7-2* and *atg5-1* mutants affect LD metabolism during dark-induced senescence, we visualized the abundance of LDs in leaf tissues by staining with the neutral-lipid-selective fluorescent dye, BODIPY493/503. An increased number of LDs was observed after 3d of darkness in WT plants (Fig. 5). However, no accumulation of LDs was observed in *atg5-1* and *atg7-2* mutants (Fig. 5). Notably, the multiple z-stacked confocal images revealed that LDs are mostly accumulated in the cytosol and not inside chloroplasts (Fig. 5A). The dynamics of LD accumulation corroborates the high levels of the plastid-derived TAG species, which were subsequently reduced after 6d of darkness in WT. Furthermore, *atg5-1* and *atg7-2* mutants displayed reduced levels of plastid-derived TAG species that can also be associated with the impairment of LD accumulation (Fig. 4A).

**Figure 5.**
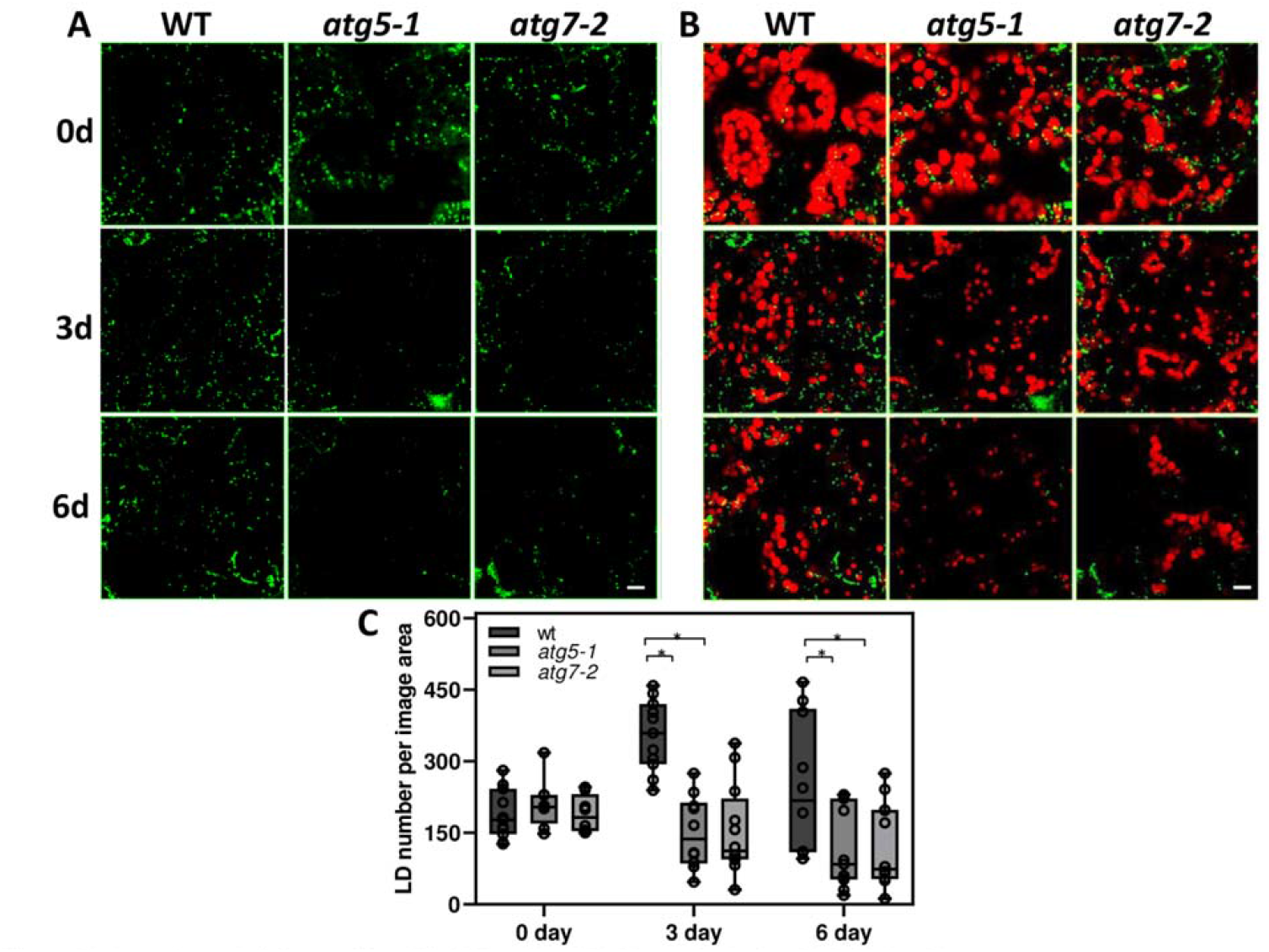
Autophagy deficiency affects Lipid Droplet (LD) dynamics during dark-induced senescence. (A) Representative confocal images of LDs (BODIPY 493/503 fluorescence, false color (green) in wild type (WT), *atg5-1*, and *atg7-2* leaves. (B) Merged image of BODIPY and chloroplast autofluorescence (false color red). Images were collected at the same magnification and are projections of Z-stacks of 20 optical sections Bar = 10 μm (C) Number of total LDs per image area. The number of LDs was quantified by ImageJ as BODIPY-stained lipid area in 2D projections of Z-stacks of 3 images of four biological replicates per genotype and time point.

### The disruption of autophagy triggers activation of PG metabolism

PG are specialized globular LDs composed by a membrane lipid monolayer surrounding a core of neutral lipids present in the chloroplast membranes (Van Wijck an Kessler, 2017). The production of PG is related to the accumulation of products from chlorophyll and chloroplast lipid catabolism such as fatty acid phytyl esters (FAPEs), TAGs and FAs (Gaude et al., 2007; Lippold et al., 2012). Given that *atg* mutants presented a substantial reduction of chloroplast lipids under extended darkness, which was not translated to an increase in TAGs or LD accumulation, we next investigated PG-related pathways by quantitative reverse transcription (RT)-PCR. The genes phytyl ester synthase 1 (*PES-1*) and *PES-2*, belonging to the esterase/lipase/thioesterase family, are required for FAPE production in Arabidopsis PG during senescence (Lippold et al., 2012). Accordingly, we observed an increase of *PES-1* and *PES-2* transcript levels following dark-induced senescence (Fig. 6A-B). However, higher induction of *PES-1* in *atg5-1* and *atg7-2*, as well as of *PES-2* in *atg7-2* mutant, was observed after 3d and 6d of darkness (Fig. 6A-B). The production of FAPE relies on both FFA released from chloroplast membranes and phytol from chlorophyll (Lippold et al., 2012). We, therefore, analyzed the chlorophyll degradation enzyme, pheophytin pheophorbide hydrolase (*PPH*). As observed for *PES1* and *PES2*, the *PPH* transcript accumulated at higher levels in *atg* mutants under dark-induced senescence in comparison to WT levels (Fig. 6D). PG serve not only as a lipid deposit but also contain an actively associated proteome (Lundquist et al., 2012). The presence of several Absence of Bc1 Complex Kinase (ABC1K) proteins was described in PG, suggesting a central role of these proteins in the regulation of PG metabolism (Ytterberg et al., 2006; Brehelin et al., 2007). The protein ABC1K7 has been characterized as a PG core component with marked accumulation during late senescence events (Lundquist et al., 2012; Bhuiyan et al., 2016). Despite the unchanged transcript levels of *ABC1K7* in WT, this gene was induced in *atg5-1* and *atg7-2* mutants following extended darkness (Fig. 6C). Although our results demonstrate a boost of PG pathways in the absence of autophagy under dark-induced senescence, *atg* mutants presented a slight but significant up-regulation for all PG-related genes even under control conditions (Fig. 6), highlighting intrinsic crosstalk between these processes.

**Figure 6.**
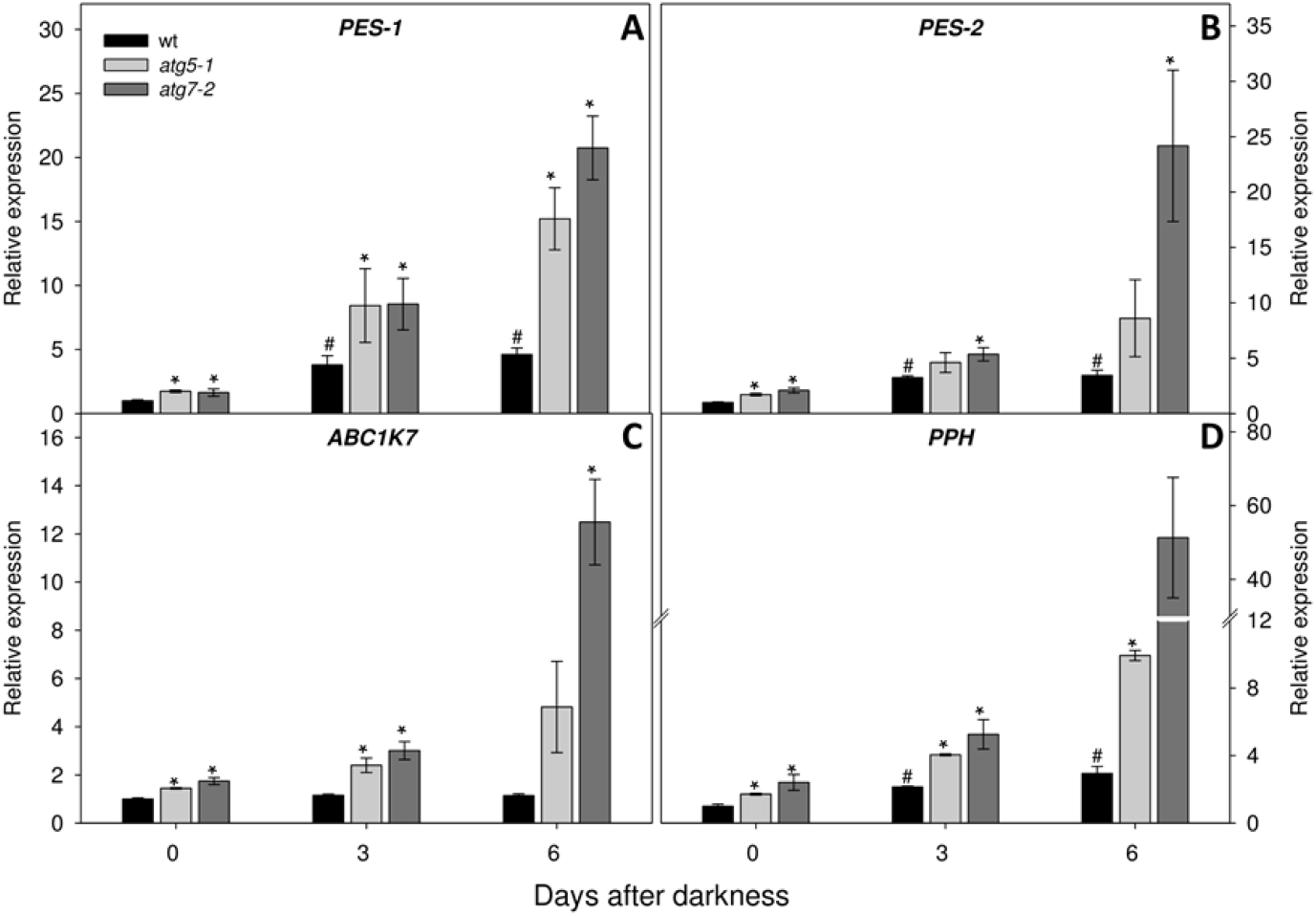
PG related genes are up-regulated in *atg* mutants during dark-induced senescence. Transcript abundance is shown for genes associated with plastoglobuli (PG), including PES-1 (A), PES-2 (B), ABC1K7 (C), PPH (D). The y-axis values represent the gene expression level relative to the wild type (WT). Data were normalized with respect to the mean response calculated for the 0-d dark-treated leaves of the WT. Values presented are means ± SE of three independent biological replicates. An asterisk (*) indicates values that were determined by Student’s t-test to be significantly different (*P* < 0.05) from the WT at each time point analyzed. A number sign (#) indicates values determined by Student’s t-test to be significantly different (*P* < 0.05) from WT at 0d.

Transmission electron microscopy analysis revealed that the number and size of PG as well as the chloroplast structure are similar in leaves of WT and *atg5-1* plants under control conditions (Figure 7A). After dark treatment, different chloroplast responses were observed in WT and *atg5-1* plants (Figure 7A). While WT presented a typical thylakoid stacking response of darkened chloroplast (Wood et al., 2019), *atg5-1* displayed smaller chloroplasts with disrupted structural remodeling, containing structures resembling senescence-associated vacuoles (Otegui et al., 2005). Moreover, a marked presence of PG was observed in *atg5-1* chloroplasts (Figure 7A). This response was ascribed to an increase in the size of PG, which reached up to 0.02 µm^2^ of area (Figure 7B-C). Otherwise, WT plants also presented increases in PG size, however, to a far less extent compared to *atg5-1* mutant (Figure 7B-C).

**Figure 7:**
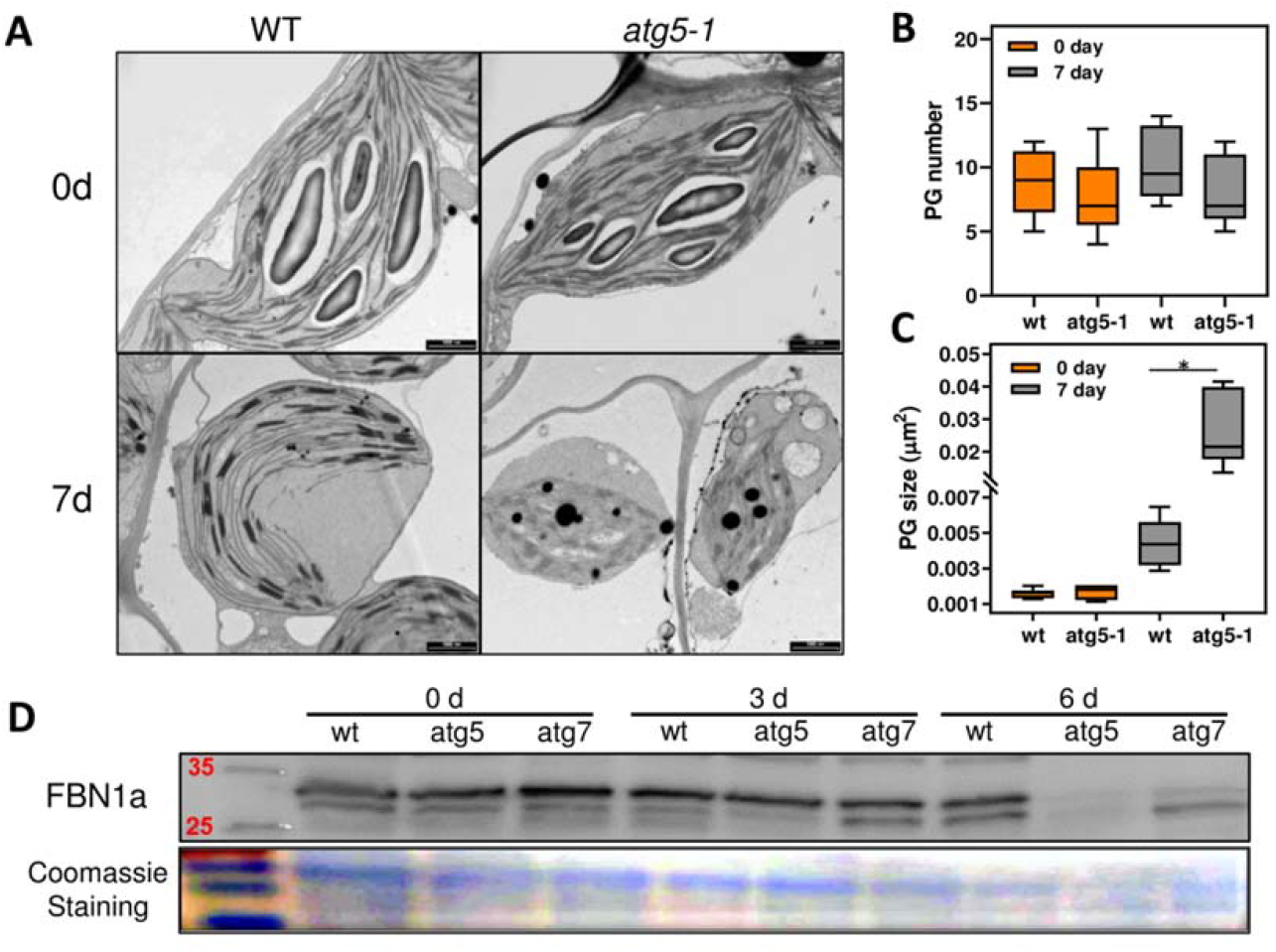
Autophagy disruption triggers altered PG response. (A) Transmission electron micrographs of leaves from 4-week-old Arabidopsis plants immediately before lights were turned off (0 d) and after further treatment for 7 days of darkness. Scale bar = 1 μm; Quantification of the number (B) and size (C) of plastoglobuli per chloroplasts (±SE, n= 6-7 chloroplasts of three independent cells); Analysis shown in A-C were performed in *atg5-1* and its respective wild type (WT). (D) Immunoblot analysis of the PG protein, FBN1a, was performed in leaves from 4-week-old Arabidopsis *atg5-1, atg7-2* and WT plants immediately before (0 d) and after further treatment for 3d and 6d in darkness. FBN1a protein display a predicted molecular mass to ∼30-35 kD. The detection of a second lower band in the region of 30kD possibly corresponds to FBN1b protein (with high sequence similarity with FBN1a) as previously observed in Giacomelli et al., 2006; Matinis et al., 2013; 2014.

Proteomic studies of PG have shown that members of the Fibrillin (FBN) family are the most abundant proteins in the PG proteome (Lundquist et al., 2012). Therefore, we next checked the content of the PG core protein FBN1a (Fibrillin 1a), previously characterized as a PG structural component (Austin 2006, Simkin et al., 2007, Singh et al., 2011). Immunoblotting analysis showed a strong reduction of FBN1a abudance in *atg5-1* mutant compared to WT, which display a slight increase of FBN1a abundance after 6d of dark treatment (Figure 7D and Supplemental Figure S2). Interestingly, the overexpression of FBN1a homologs resulted in increased numbers and clustering of PGs in tobacco leaves and tomato fruit (Rey et al., 2000; Simkin et al., 2007). Additionally, FBN overexpression was also associated with the delay of thylakoid disintegration, suggesting its role on protection/maintenance of chloroplast membranes (Simkin et al., 2007; Rottet al., 2015). The reduction of FBN1a protein abundance is most likely related to the occurrence of supersized PG coupled with the overall degradation of chloroplast membranes in *atg5-1* mutant plants under dark-induced senescence. Collectively, our findings highlight PG as a potential lipid storage site in *atg* mutants submitted to extended darkness conditions. It will therefore be interesting to determine hereafter whether and which extent FBNs determine PG structural modeling during low energy conditions.

### *atg* Mutants induce lipid oxidation pathways in response to dark-induced senescence

FA oxidation may be of particular significance when the carbon and energy status is rather low. During carbon starvation, the importance of lipid metabolism was demonstrated by the characterization of β-oxidation loss-of-function mutants (Kunz et al., 2009). In addition, autophagy association with peroxisomal protein abundance was recently demonstrated (McLoughlin et al., 2018, 2020; Have et al., 2019). The general reduction of chloroplast lipids, together with reduced LD numbers in *atg*5*-1* and *atg*7-*2* mutants subjected to extended darkness, raised the question of whether these lipids were being quickly oxidized to be used as energetic substrates in *atg* mutants. We, therefore, evaluated the expression of genes related to LD catabolism and peroxisomal pathways of FA oxidation. We observed that the transcript levels of the peroxisomal enzymes Acyl-CoA oxidase (*ACX-4*), Multifunctional protein 2 (*MPF-2*), Ketoacyl-CoA thiolase (*KAT-2*), Citrate synthase 1 (*CSY-*1) and peroxisomal Malate Dehydrogenase (*pMDH1*) genes were generally higher in *atg5-1* and *atg7-2* under extended darkness (Fig. 8 A-E). More specifically, *ACX-4* and *CSY-1* were slightly induced in WT throughout the treatment. However, higher transcript levels were observed in *atg5-1* and *atg7-2* plants after 6d of darkness (Fig. 8A-B). Despite no major changes in *MPF-2* and *KAT-2* transcript abundance in WT plants, trends of increase were observed in *atg5-1* and *atg7-2* after 9d of darkness (Fig. 8C-D). We further confirmed that *pMDH-1* was downregulated following dark-induced senescence. *pMDH-1* operates in the reduction of oxaloacetate to malate and enables the re-oxidation of NADH for continued β-oxidation (Pracharoenwattana et al., 2007). Given that *pMDH-1* is not participating directly in β-oxidation, it is possible that an alternative pathway of NADH oxidation becomes operational under carbon starvation.

**Figure 8.**
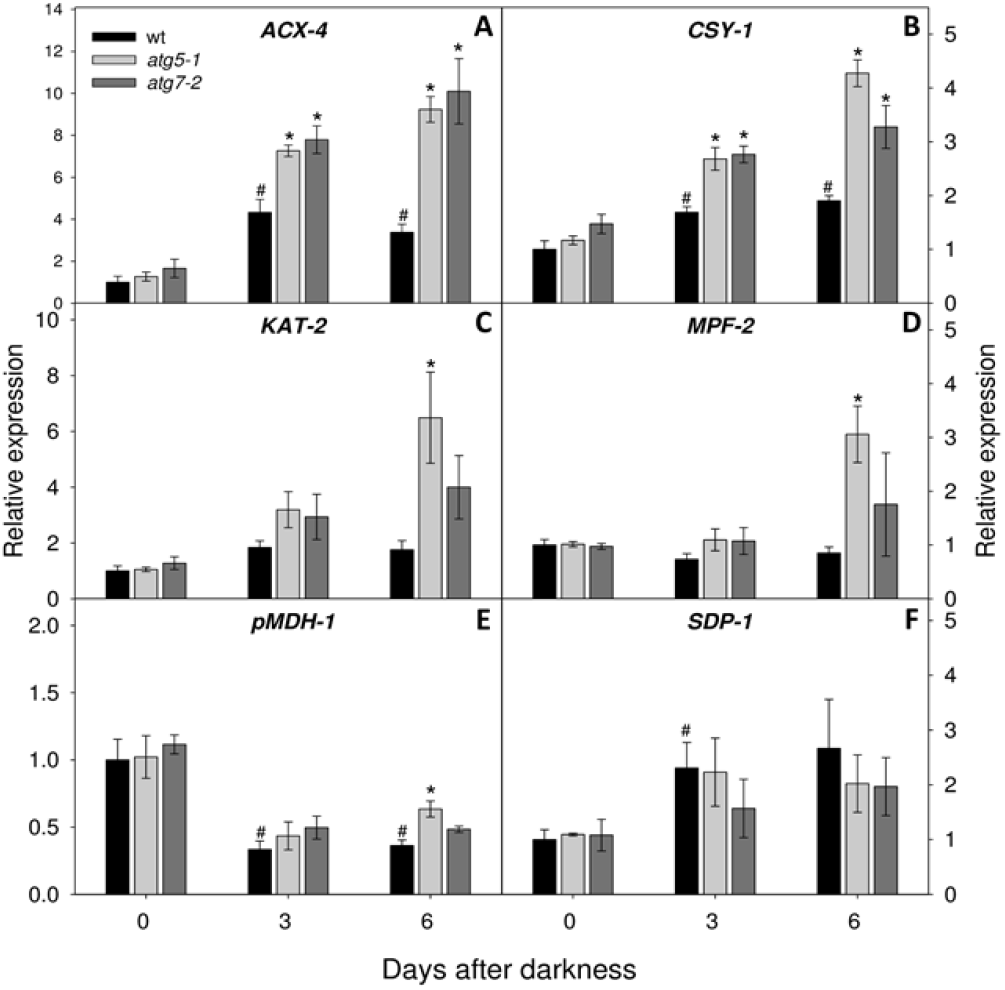
Changes in transcript levels in WT and *atg* mutants during dark-induced senescence. Transcript abundance is shown for genes associated with lipid catabolic pathways, including ACX-4 (A), CSY-1(B) KAT-2 (C), MPF-2 (D), pMDH-1 (E), SDP-1 (F). The y-axis values represent the expression level relative to the wild type (WT). Data were normalized with respect to the mean response calculated for the 0-d dark-treated leaves of the WT. Values presented are means ± SE of three independent biological replicates. An asterisk (*) indicates values that were determined by Student’s t-test to be significantly different (*P* < 0.05) from the WT at each time point analyzed. A number sign (#) indicates values determined by Student’s t-test to be significantly different (*P* < 0.05) from WT at 0d.

To investigate whether the reduced number of LDs is a result of the induction of catabolic pathways or impairment in biosynthesis, we next evaluated the expression of *SDP-1* gene. Despite being characterized as the primary lipase mediating LD catabolism in seeds, compelling evidence has suggested that *SDP-1* also operates in senescent and carbon-starved leaves (Fan et al., 2017, 2019). Indeed, an accumulation of *SDP-1* transcript levels was observed in WT plants during the early stages of darkness (Fig. 8F). Nonetheless, no significant difference in *SDP-1* transcript levels between WT plants and *atg5-1* and *atg7-2* was observed, suggesting that the reduced levels of LDs are likely more related to the impairment of LD biogenesis rather than an induction of LD degradation pathways.

### Altered FFA response of *atg* mutants under dark-induced senescence

Since autophagy deficiency altered cytosolic LD dynamics and β-oxidation pathways, we next evaluated if this response also affected the pool of free FA (FFA). Figure 9 shows that long-chain FFA (between 13C-21C chain length) are the more affected by both genotype and darkness, while no major changes were observed for very long-chain FFA (> 22C). Higher levels of 16:3, 18:1, 18:2, 18:3 in *atg5-1* and *atg7-2* genotypes under control conditions were observed, reinforcing the functional role of autophagy in basal lipid homeostasis. Upon darkness treatment, the levels of C18 and C20 FFAs were reduced, and unsaturated C16 accumulated in *atg5-1* and *atg7-2* mutants (Fig. 9). By contrast, WT plants presented an initial increase followed by later reduction of C16 and C18 FFAs except for 16:3 and 18:1 species FFAs, which were respectively, accumulated and reduced throughout darkness (Fig. 9). Although unsaturated C16 and C18 FAs are typical components of chloroplast membranes, only 16:3 is exclusively found in the thylakoid lipid MGDG in plant species (e.g. Arabidopsis), and it is, therefore, a chloroplast lipid marker (Ohlrogge et al., 1991; Kunz et al., 2009; Hölzl and Dörmann, 2019). Thus, the overall accumulation of 16:3 is in good agreement with the reduction of galactolipids observed under dark-induced senescence. Additionally, this result suggests that the rate of 16:3 release exceeds its channeling to LD and PG. The higher accumulation of 16:3 in *atg* mutants strengthens our claim that this disbalance is more intense when autophagy is disrupted.

**Figure 9.**
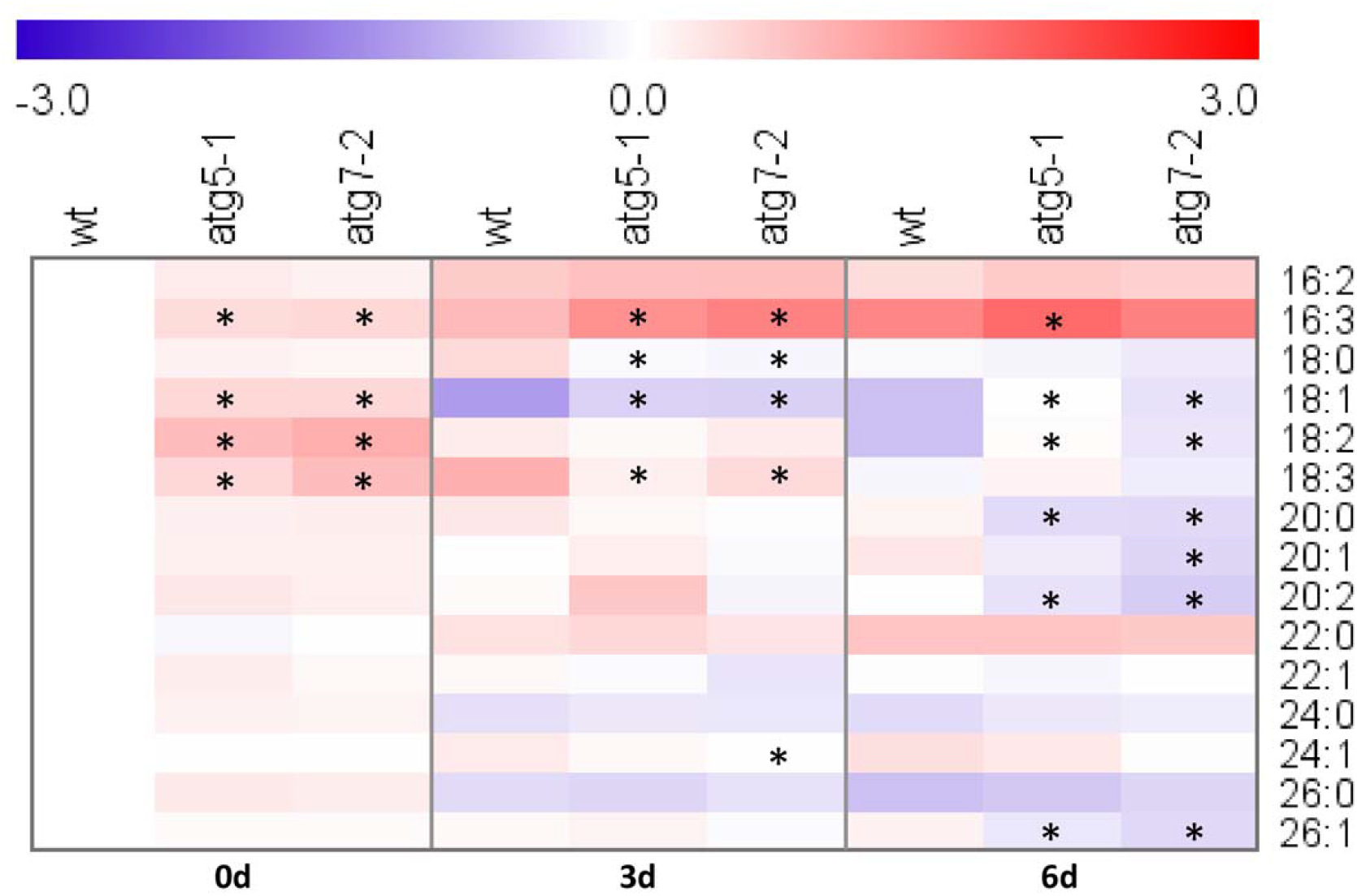
Changes in free Fatty Acid levels in WT and *atg* mutant plants under dark-induced senescence. Values plotted are log_2_ fold change. Data were normalized to the mean response calculated for the 0-d dark treated leaves of the wild-type (WT). Values presented are means ± SE of five-six biological replicates per genotype; an asterisk (*) designate values that were determined by the Student’s t-test to be significantly different (*P* < 0.05) from WT each time point analyzed.

## Discussion

It is well established that plant autophagy plays a crucial role in the energetic metabolism mediating the supply of proteins and amino acids under carbon starvation conditions (Avin-Wittenberg et al., 2015; Barros et al., 2017; Hirota et al., 2018; McLoughlin et al., 2020). Furthermore, in both animals and yeast, autophagy is recognized as an important lipid catabolic pathway in starved cells (Singh et al., 2009). In plant leaves, growing evidence has highlighted the role of autophagy in managing cell lipid components and reserves (Avin-Wittenberg et al., 2015; McLoughlin et al., 2018,2020; Fan et al., 2019; Have et al., 2019). In fact, the lipid response of *atg* mutants under N starvation has been previously demonstrated (McLoughlin et al., 2018; Have et al., 2019). More recently, McLoughlin et al. (2020) reported extensive lipid alterations in maize *atg12* mutants submitted to carbon deprivation induced by short-term darkness. Here, we provide further evidence that autophagy broadly affects lipid and LD metabolism during dark-induced senescence in Arabidopsis. Notably, similarities and differences between the aforementioned studies and the findings reported here were observed. The main points were summarized in Supplemental Figure S3, and they will be further discussed throughout the text.

By performing a detailed lipidomics analysis, we revealed that autophagy widely affects the lipid homeostasis of Arabidopsis during dark-induced senescence. The major differences reported here are related to the *atg5-1* and *atg7-2* mutant plants, characterized by a full inhibition of autophagy (Chung et al., 2010). By contrast, *atg9-4*, which is characterized by a partial reduction of autophagy, presented an intermediary impact on lipid metabolism that was more evident during the late stages of extended darkness (Fig. 1 and Supplemental Figure S1). The fact that *atg9-4* did not cluster with *atg5-1* and *atg7-2* implies that the lipid response is directly related to autophagy intensity. Indeed, the characterization of Arabidopsis *ATG*5 and *ATG*7 knockout and overexpression mutants showed a positive correlation between the level of autophagy and seed FA content (Minina et al., 2018). We previously demonstrated that amino acid catabolic pathways were less compromised in *atg9-4* mutants, contributing to its more tolerant phenotype during dark-induced senescence (Barros et al., 2017). Additionally, the limited reproductive phenotype of *atg* mutants is also dependent on autophagy intensity since parameters related to seed and silique production were more compromised in *atg* lines fully defective in autophagy (Barros et al., 2017). Taking into account the fact that Arabidopsis is an oilseed (Li et al., 2006), the results reported here and by others (Minina et al., 2018) suggest that the altered reproductive and carbon starved phenotype of *atg* mutants is likely not only a consequence of impairment of protein catabolic process but is also derived from an extensive reprogramming of lipid metabolism.

The detailed analysis of the specific lipid classes revealed an altered lipid phenotype in mutants lacking autophagy under non-stress conditions. These differences are mostly attributed to galactolipids and the phospholipids PC and PE, whereas minor differences were observed for TAGs (Figs 2, 3, and 4). The galactolipids MGDG and DGDG are prevalent in the chloroplast envelope and the thylakoid membrane (Hölzl and Dörmann, 2019). By contrast, PC and PE are highly abundant in the plasma membrane, mitochondrial membrane, and endoplasmic reticulum (Liu et al., 2015). Hence, the high levels of PC and low levels of MGDG and DGDG reveal an imbalance of cytosolic and plastidial membrane lipids in *atg5-1* and *atg7-2* mutants under normal growth conditions. Similar results were previously reported in Arabidopsis *atg5* mutants, which displayed a general alteration of phospholipid levels under non-stressful conditions (Have et al., 2019). The accumulation of lipid-breakdown products in maize *atg12* mutants also suggested an induced turnover of membrane components under normal growth conditions (McLoughlin et al., 2018, 2020). In addition, Fan et al., 2019 reported that the disrupted membrane lipid turnover in *atg* mutants affected TAG synthesis. The TAG profile detailed here reveals that only specific TAG species are reduced in *atg5-1*, and *atg7-2* mutants under normal growth conditions, which seems to contribute to the low total TAG content previously reported.

Collectively, our results support the notion that, beyond the role of autophagy in basal lipid metabolism, autophagy is clearly required for lipid remodeling under starvation conditions in Arabidopsis. Upon exposure to extended darkness, PC metabolism is broadly rearranged, presenting different patterns of accumulation and reduction (Fig. 2). In general, the majority of 32C and 34C PC species increases, whereas 36C and 38C declines upon darkness (Fig. 2). However, an altered response of PC accumulation and degradation was observed in *atg* mutants. The turnover of membrane lipids is a recognized aspect of plant senescence in general (Troncoso-Ponce et al., 2013). It seems that the disruption of autophagy leads to a differential degradation rate of membrane lipid components, most likely affecting the supply of FAs to sustain other cellular processes, including respiration. Alteration of PE levels was also observed in *atg* mutants under our experimental conditions (Fig. 2). Noteworthy, PE is an essential component of ATG8 conjugation during autophagosome membrane biogenesis (Soto-Burgo et al., 2018; Barros et al., 2020). The less accentuated reductions and accumulation of several PE species (PE 36:4, 36:3(1), 38:2, 36:5, 38:3) in *atg* mutants might be related to the disrupted usage and recycling of these molecules for autophagosome biogenesis, as previously suggested (Avin-Wittenberg et al., 2015; Have et al., 2019).

Apart from changes in the extra-plastidial phospholipids, *atg7-2* and *atg5-1* mutants also presented intense degradation of chloroplastic MGDG and DGDG during dark-induced senescence (Fig. 5). This data is consistent with the hypersensitive phenotype marked by chloroplast dismantling observed in *atg* mutants submitted to dark-induced senescence (Thompson et al., 2005; Phillips et al., 2008; Barros et al., 2017). Accordingly, autophagy has been characterized to mediate the degradation of chloroplast proteins by different selective processes (Ishida et al., 2008; Wada et al., 2009; Michaeli et al., 2014; Izumi et al., 2017; Nakamura et al., 2018). This fact aside, the results shown here reinforce the hypothesis that other pathways likely mediate the degradation of chloroplast lipid components in carbon-starved plants and that they are likely to be more active when autophagy is absent. Moreover, this response is not restricted to carbon limitation, as similar results were reported in both Arabidopsis and maize *atg* mutants under low-nitrogen conditions (Supplemental Figure S3) (McLoughlin et al., 2018; Have et al., 2019).

The breakdown of membrane lipid components releases FFAs for further oxidation in peroxisomes. The overaccumulation of FFAs is detrimental to the plant cell. Consequently, FFAs are first incorporated as neutral lipids, to then be directed to peroxisomal FA catabolic pathways (Fan et al., 2017). TAGs stored in LDs are not abundant in leaf mesophyll cells. However, their number substantially increases under abiotic conditions and developmental senescence (Pyc et al., 2017; Yang et al., 2018). In agreement with this, we observed a transient LD accumulation after 3d of darkness in WT plants. In addition, a marked increase of highly unsaturated TAG levels was observed after 3d of darkness in WT (Fig. 3). Despite an increase of some FFA species after 3d of darkness, this increase was not overloaded to result in a lipotoxic effect since WT plants present favorable performance under dark-induced senescence (Barros et al., 2017). Indeed, the pattern of initial accumulation and later reduction of FFA 16:2, 18:0, 18:2, 18:3 was similar to the observed for TAG and LD (Figs. 4, 5 and 9), suggesting a dynamic equilibrium between membrane consumption, lipid reserve formation, and FA utilization in WT plants. By contrast, not only LD accumulation was disrupted in *atg5-1* and *atg7-2* mutants, but they also showed lower levels of unsaturated TAGs and higher accumulation of plastidial FFA 16:3 after 3d and 6d of darkness. Collectively, these results demonstrate that autophagy is required for the proper mobilization of chloroplast lipids destined for cytosolic LD biosynthesis in Arabidopsis starved cells. A recent study of maize *atg12* mutants subjected to carbon deprivation reported distinct LD dynamics (McLoughlin et al., 2020). It was observed that *atg12* mutants contained more cytoplasmic LD under normal growth conditions. In individually darkened leaves, LD numbers were not affected in WT and only slightly reduced in the *atg12* mutant. A possible explanation for this discrepancy may stem from the experimental systems utilized. While McLoughlin et al. (2020) analyzed individually darkened leaves of maize, we examined Arabidopsis in a whole plant darkening system. Plants have divergent metabolic strategies in response to different darkening treatments (Law et al., 2018). It has been postulated that individually darkened leaves still receive sugars from other parts of the plant during short-term dark periods. However, the lipid catabolic response is not activated as long as carbohydrates are available, which likely affects LD dynamics (Kunz et al., 2009; Law et al., 2018). Not only LD responses diverge according to the darkness treatment imposed, but the results reported until now also open new avenues regarding contrasting LD responses in different plant species.

In Arabidopsis, autophagy operates in the mobilization of FAs from organelle membranes to TAG synthesis under normal growth conditions (Fan et al., 2019). The same study stated that autophagy has no influence on TAG formation during darkness, but instead operates in LD degradation by lipophagy mechanism. Despite that, general TAG accumulation was observed in *atg* mutants after 6d of darkness, which was not accompanied by cytosolic LD accumulation (Figs 4 and 5). Nonetheless, caution must be taken when comparing the results obtained here with previous findings (Fan et al., 2019) due to the differences of the genotypes analyzed (Barros et al., 2020). Thus, although Fan et al. (2019) demonstrated the existence of lipophagy in different mutant backgrounds, we analyzed plants solely lacking autophagy. When considered together, our results, coupled with the results from Fan and colleagues, highlight the potential role of autophagy as a versatile player in LD metabolism, particularly in starved cells.

Beyond the LDs in the cytosol, PG serve as subcompartments for lipid biosynthesis and storage in chloroplasts (Rottet et al., 2015; van Wijk and Kessler, 2017). The analysis of PG-related genes indicates a marked activation of PG metabolism in *atg5-1* and *atg7-2* mutants during dark-induced senescence (Fig. 6). In agreement with that, we observed the presence of large PG in chloroplasts of *atg5-1* mutants under extended darkness conditions (Fig. 7A-C). On the other hand, a reduced abundance of the PG core protein FBN1a was observed in *atg* mutants after 6d of darkness (Fig. 7D and Supplemental Figure S2). It is noteworthy that fibrillin proteins, localized on the PG surface, were proposed to play both structural and protection roles in chloroplast, preventing the PG coalescence and delaying thylakoid disintegration (Austin et al., 2006; Ytterberg et al., 2006; Simkin et al., 2007; Singh et al., 2011; Rottet al., 2015). It is tempting to suggest, therefore, that the reduction of FBN1a protein can be associated with the large size of PG as well as chloroplast sensitive phenotype in *atg* mutants in response to dark-induced senescence.There is a strong correlation between thylakoid dismantling and supersizing of PG during senescence, which is likely associated with the massive deposit of FAs and phytol, released from chloroplast degradation (Lippold et al., 2012; Besagni et al., 2013; Rottet et al., 2015; van Wijk and Kessler, 2017). In light of this, we also verified an increase of *PPH* transcript in *atg5-1* and *atg7-2* mutants compared to WT under darkness (Fig. 6D). It seems reasonable to assume that both the products from the degradation of chlorophyll (Barros et al., 2017) and chloroplast membranes are funnelled for PG formation in *atg* mutants submitted to extended darkness (Fig. 3,7,10). This response can be presumed as a strategic mechanism preventing the accumulation of lipotoxic intermediaries. Collectively, our results imply that the disruption of autophagy triggers a differential response prompting PG rather than cytosolic LD formation under dark-induced senescence. Nonetheless, whether PG can be remobilized to provide lipid-derived energetic substrates remains to be further investigated.

**Figure 10.**
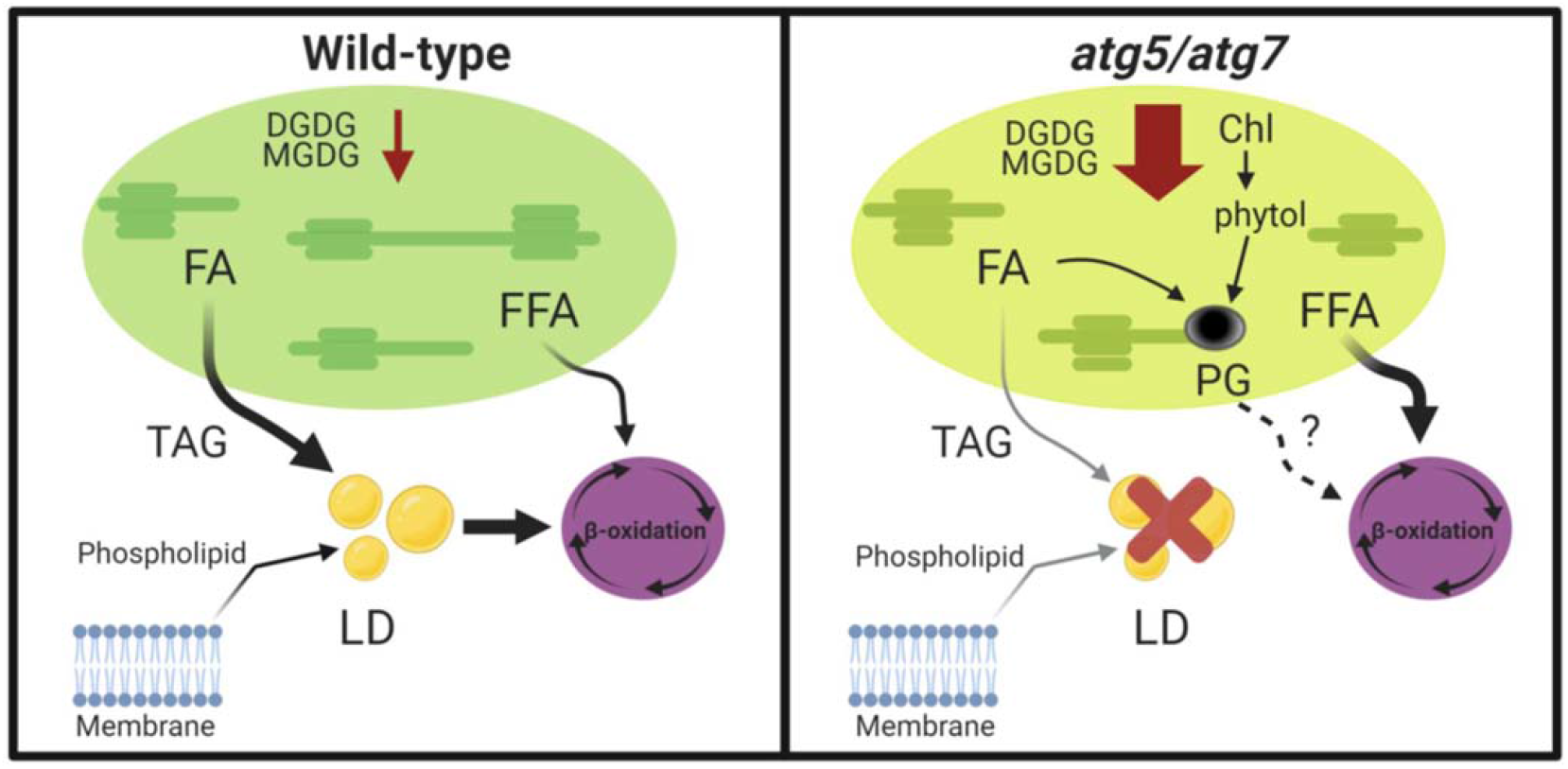
Schematic model of lipid remodeling response of *atg* mutants during extended darkness. Under extended darkness, the plastidial and extraplastidial membranes, composed by MGDG/DGDG and phospholipids, respectively, are degraded, releasing FAs to be stored as TAGs in LD in the cytosol. LDs and transient FFA donate substrates for β-oxidation in peroxisomes. Meanwhile, in *atg* mutants, there is a substantial degradation of chloroplast lipids, while the production of cytosolic LDs is impaired. Part of the released plastidial FAs, together with phytol derived from chlorophyll degradation, are channeled to the production of PG in the chloroplast. β-oxidation pathways are highly induced to consume overloaded plastidial FFA and, possibly, PG lipids (question mark). Abbreviations: Digalactosyldiacylglycerol (DGDG); Fatty acids (FA); Free Fatty Acids (FFA); Lipid Droplet (LD); Monogalactosyldiacylglycerol (MGDG).

It is already known that, during extended darkness, FAs are oxidized by β-oxidation, generating acetyl-CoA that, combined with oxaloacetate, produces citrate to be exported to mitochondria for ATP generation (Kunz et al., 2009). To further understand whether the released membrane lipid components were channeled to the maintenance of the energetic status, we paid particular attention to the transcriptional activation of lipid oxidation pathways. Our results show an up-regulation of peroxisomal genes in plants with disrupted autophagy after 6d and 9d of darkness (Fig. 8). Notably, the transcriptional induction of β-oxidation has been correlated to mitochondrial activity in darkened leaves (Law et al., 2018). Interestingly, the same *atg* mutants exhibited higher rates of respiration and an elegant metabolic reprogramming under carbon starvation conditions (Avin-Winttenberg et al., 2015; Barros et al., 2017).

Autophagy also plays a role in peroxisome quality control by the selective degradation of damaged peroxisomes, a process known as pexophagy (Marshall and Vierstra, 2018). The accumulation of peroxisome-related transcripts/proteins in *atg* mutants has been reported lately, and it can represent both activation of peroxisomal metabolism or accumulation of oxidized peroxisome (Supplemental Figure S3) (McLoughlin et al., 2018, 2020; Have et al.,2019). Specifically, under carbon starvation conditions, the enrichment of peroxisomal proteins was followed by an increase of glyoxysomal proteins (McLoughlin et al., 2020). This response suggests that peroxisomes are most likely converted to glyoxysomes to sustain high activation of FA ß-oxidation pathways during starvation conditions (McLoughlin et al., 2020). The induction of peroxisomal pathways reported here is positively correlated with the high levels of plastidial FFAs in *atg* mutants, which can be interpreted as a response to control FFA levels concomitantly with energy production. When taken together with our work, these studies suggest that the primary metabolism of *atg* mutants is not only rearranged for amino acid catabolism but is also possibly involved in supporting lipid oxidation pathways. In light of this, further studies are clearly required (*i*) to investigate whether the disruption of LD dynamics and differential membrane recycling of *atg* mutants impact the supply of lipid substrates for FA oxidation pathways and (*ii*) to characterize the function of autophagy under a range of environmental stresses during this process.

## Conclusions

In the current work, we present compelling evidence regarding the importance of autophagy to lipid homeostasis in Arabidopsis leaves. During carbon starvation, the products of organellar membrane recycling are channeled to cytosolic LD production. This enables the balance of FA production and turnover. The functional lack of autophagy impacts proper membrane mobilization and activates a general chloroplast lipid degradation program while failing to produce cytosolic LDs. Nonetheless, the products of chloroplast membrane dismantling are possibly channeled to PG generation in *atg* mutants chloroplasts (Fig. 10). We postulate that the increased activation of peroxisomal lipid catabolic pathways is part of a complex metabolic response to catabolize the FFAs released from chloroplast dismantling, and possibly, from PG mobilization. Although our results highlight novel insights regarding the role of autophagy in lipid metabolism, they certainly underestimate the complex crosstalk between autophagy, peroxisome, mitochondria, and the endoplasmic reticulum. An investigation of the interplay involving autophagy, membrane recycling, and organellar maintenance is required to fully understand the implications of autophagy for the regulation of lipid reserve dynamics and energy homeostasis. The combined results presented here are of potential application for the breeding of crops with a higher yield, increased resilience, and it seems reasonable to assume that there is a timely need to accelerate our understanding of the mechanisms controlling autophagy and associated processes in response to environmental conditions.

## Methods

### Plant Material and Dark Treatment

All *Arabidopsis thaliana* (L.) Heynh plants used in this study were of the Columbia ecotype (Col-0). The T-DNA mutant lines *atg9-4* (SALK_145980) (Shin et al., 2014), *atg5-1* (SAIL_129B079) (Yoshimoto et al., 2009), *atg7-2* (GK-655B06) (Hofius et al., 2009) were used in this study. Seeds were surface-sterilized and imbibed for 4 days at 4°C in the dark on 0.7% (w/v) agar plates containing half-strength Murashige and Skoog (MS) media (Sigma-Aldrich; pH 5.7). Seeds were subsequently germinated and grown at 22°C under short-day conditions (8 h light/16 h dark), 60% relative humidity with 150 μmol photons m^-2^ s^-1^. For dark treatments and subsequent induction of dark-induced senescence, 10- to 14-day-old seedlings were transferred to soil and then grown at 22°C under short-day conditions (8 h light/16 h dark). Four-week-old plants were transferred to darkness in the same growth cabinet. Whole rosettes of two different plants representing an independent replicated by genotype were harvested at intervals of 0, 3, 6, 9 days after transition to darkness. At least five biological replicates of each genotype were obtained. They were immediately frozen in liquid nitrogen and stored at −80^°^C until further analysis.

### Visualization and Analysis of LDs

To visualize LDs in Arabidopsis leaves, LDs were stained with BODIPY 493/503 (Invitrogen; from 4 mg/mL stock in DMSO). The sixth rosette leaf of 4-week-old plants was used for LD visualization and analysis. Leaf discs were stained with BODIPY solution (2μg/mL) in the dark and under vacuum for 10 min. Confocal images were acquired using the FV-1200 confocal microscope (Olympus, Japan) with a 60X/1.42 oil immersion objective. Excitation was performed at 488nm, and emission was collected between 500-540nm for BODIPY 493/503 and 650-750nm for the chlorophyll autofluorescence. The LD number was quantified using ImageJ (Schneider et al., 2012). Z-stack series of three images from four plants for each genotype were used for the quantification of LDs.

### Lipid Profiling and Free Fatty Acid Analysis

Lipids and natural endogenic free fatty acids were extracted from six biological replicates using the MTBE method described by Salem et al. (2016). Vacuum-dried organic phases were processed using ultra-performance liquid chromatography (on a C8 reverse-phase column) coupled with Fourier transform mass spectrometry (exactive mass spectrometer; Thermo Fisher Scientific) in positive and negative ionization modes. The free fatty acid were analysed using ultra-performance liquid chromatography (Waters, Acquity I Class System) coupled with mass spectrometer (Q-Exactive Thermo Fisher Scientific, H-ESI). Processing of chromatograms, peak detection, and integration were performed using REFINER MSH 10 (GeneData). Processing of mass spectrometry data included the removal of the fragmentation information, isotopic peaks, and chemical noise. Selected features were annotated using an in-house lipid database (Lapidot-Cohen et al., 2020). Five-six biological replicates were used for the analysis.

### Expression analysis by RT-PCR

Total RNA was isolated using TRIzol reagent (Ambion, Life Technology) according to the manufacturer’s recommendations. The total RNA was treated with TURBO DNA-free kit (Invitrogen). The integrity of the RNA was checked on 1% (w/v) agarose gels, and the concentration was measured using a Nanodrop spectrophotometer. Finally, 1 μg of total RNA was reverse transcribed with SensiFAST cDNA Synthesis Kit (Bioline) according to the manufacturer’s recommendations. Real-time PCR reactions were performed in a 384-well microlitre using SensiFAST SYBR® No-ROX Kit (Bioline). The primers used here were designed using the open-source program QuantPrime-qPCR primer designed tool (Arvidsson et al., 2008) and are described in Table S2. The transcript abundance was calculated by the standard curves of each selected gene and normalized using the constitutively expressed genes ACTIN (AT2G37620). Three biological replicates were processed for each experimental condition.

### Transmission Electron Microscopy

Small pieces of leaves were cut and fixed in 3% Glutaraldeyde in 0.1M Cacodylate buffer (pH 7.4) 10 hours at room temperature in a desiccator and then transferred to 40°C for the continuation of fixation overnight. The tissues were washed in cacodylate buffer and postfixed and stained with 2% osmium tetroxide, 1.5% potassium ferricyanide in 0.1M cacodylate buffer for 2 hours. Tissues were then washed in cacodylate buffer and dehydrated through a graded series of ethanol treatments for 10 min each step, followed by 100% anhydrous ethanol 3 times, 20 min each, and propylene oxide 2 times, 10 min each. The tissues were infiltrated with increasing concentrations of Agar 100 resin in propylene oxide for 16 hours each step. The tissues were embedded in fresh resin in a 600°C oven for 48 hours. Ultrathin sections, approximately 80 nm thick, were cut on a Leica Reichert Ultracut S microtome, collected onto 200 Mesh carbon-formvar coated copper grids, and stained with Uranyl acetate followed by Reynold’s Lead Citrate for 10 min. The sections were examined using Tecnai 12 TEM 100kV (Phillips, Eindhoven, the Netherlands) equipped with MegaView II CCD camera and Analysis® version 3.0 software (SoftImaging System GmbH, Münstar, Germany).

### Immunoblot Analyses

Plant leaf tissue (∼100 mg) was ground in liquid nitrogen, and total proteins were extracted using a protein extraction buffer previously described (Gruis et al., 2002). After the addition of buffer, centrifuged for 15 min at 20,000g; supernatants were collected, and samples were incubated immediately for 10 min at 100°C, vortexed, and samples were adjusted in SDS-PAGE sample buffer (Laemmli, 1970). Total proteins were separated by SDS-PAGE and transferred to a polyvinylidene difluoride membrane (GE Healthcare Life science). Equal loading was validated using Coomassie Brilliant Blue staining of the membrane. For immunodetection, antibodies raised against FBN1a (Agrisera; AS06116) was used in combination with horseradish peroxidase-conjugated goat anti-rabbit antibody (Sigma; ab920)

### Statistical analyses

The experiments were conducted in a completely randomized design with 3-6 replicates of each genotype. Data were statistically examined using analysis of variance and tested for significant (*P* < 0.05) differences using Student’s *t* tests from algorithm embedded into Microsoft Excel. Principal component analysis (PCA) was performed to reduce the dataset and identify the variables that best explained the highest proportion of total variance using Metaboanalyst 4.0, an open-source web-based tool for metabolomics data analysis (Chong et al., 2019).

### Accession Numbers

The Arabidopsis Genome Initiative locus numbers for the main genes discussed in this article are as follows: *ATG5* (At5g17290), *ATG7* (At5g45900), and *ATG9* (At2g31260).

## Supplemental Data

The following supplemental materials are available.

**Supplemental Figure S1:** Lipid response of *atg* mutants throughout dark-induced senescence. Principal component analysis (PCA) was performed using the lipid data of *atg5-1, atg7-2, atg9-4* and wild type (WT) genotypes after 0 (0d), 3 (3d), (6d), and 9 (9d) days of dark treatment.

**Supplemental Figure S2: Quantification of the FBN1a protein signal**

Integrated density values for FBN1a were first normalized to loading control and then expressed as a percentage of the obtained value for the wild type (WT) at 0d. Data represent means ±SE; n= 3 biological replicates. An asterisk (*) indicates values that were determined by Student’s t-test to be significantly different (*P* < 0.05) from the WT at each time point analyzed.

**Supplemental Figure S3:** Compared lipidomics of Arabidopsis and maize *atg* mutants under carbon and nitrogen starvation. General representation of the changes in Phospholipids, Galactolipds, Lipid Droplet number, and β-oxidation transcripts reported in Arabidopsis and maize under carbon starvation (this study; McLoughlin et al., 2020) and nitrogen starvation (McLoughlin et al., 2018; Have et al., 2019). The scheme does not take into consideration the experimental differences in these studies. Abbreviations-PC: Phosphatidylcholine; PE-: Phosphatidylethanolamine; PG: phosphatidylglycerol; - C: Carbon starvation; -N: Nitrogen starvation

**Supplemental Table S1**: Summary of the main variables of the first three Principal Component (PC) of *atg* mutants and wild type (WT) genotypes after 0 and 6 days of dark treatment. The contribution of which PC is showed as cumulative variance (CV).

**Supplemental Table S2:** Primers used in the RT-PCR analyses performed in this study **Supplemental Table S3:** Relative lipid content of leaves of Arabidopsis knockout mutants *atg5-1, atg7-2, atg9-4*, and wild type (WT) after 0, 3, 6, and 9 days of dark treatment

**Supplemental Table S4:** Variables of the first three Principal Component (PC) of *atg* mutants and wild type (WT) genotypes after 0 and 6 days of dark treatment

## ACKNOWLEDGEMENTS

We thank Prof. Oren Ostersetzer-Biran from The Hebrew University of Jerusalem for providing the facilities allowing transcript analysis, as well as Corinne Best and Sofia Shevtsov, for their assistance with qPCR analyses. We also thank Dr. Naomi Melamed-Book and Dr. Yael Friedmann from the microscopy core facility at The Hebrew University’s Life Science core facility for her technical support with confocal and EM analysis. We also thank Prof. Zack Adam from The Hebrew University of Jerusalem for providing us with the anti-FBN1a antibody.

