## Supplemental figures and data S1,S2 for "Autophagy is required for lipid homeostasis during dark-induced senescence in Arabidopsis"

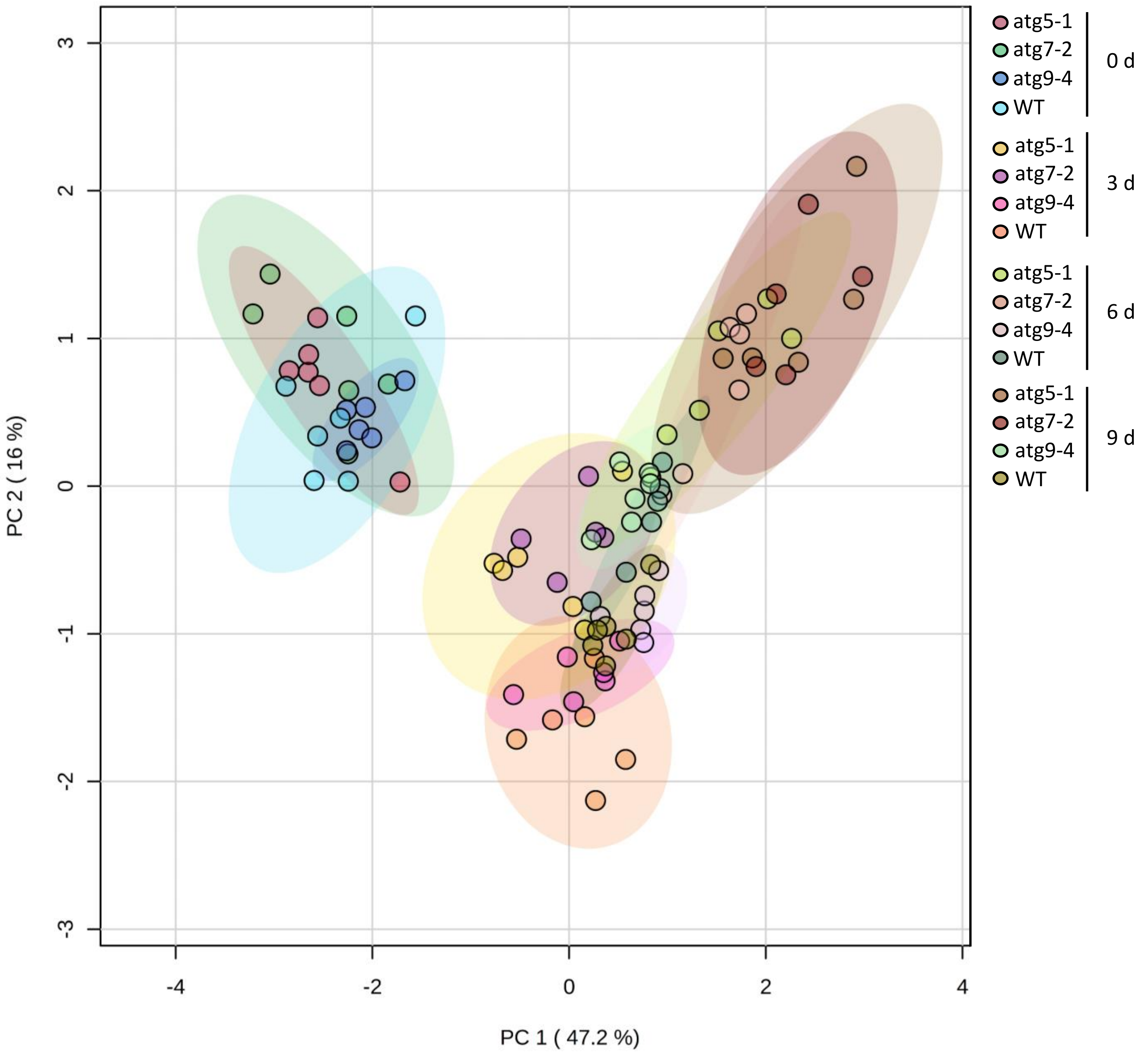


**Supplemental Figure S1:** Lipid response of *atg* mutants throughout dark-induced senescence. Principal component analysis (PCA) was performed using the lipid data of *atg5-1*, *atg7-2*, *atg9-4* and wild type (WT) genotypes after 0 (0d), 3 (3d), (6d), and 9 (9d) days of dark treatment

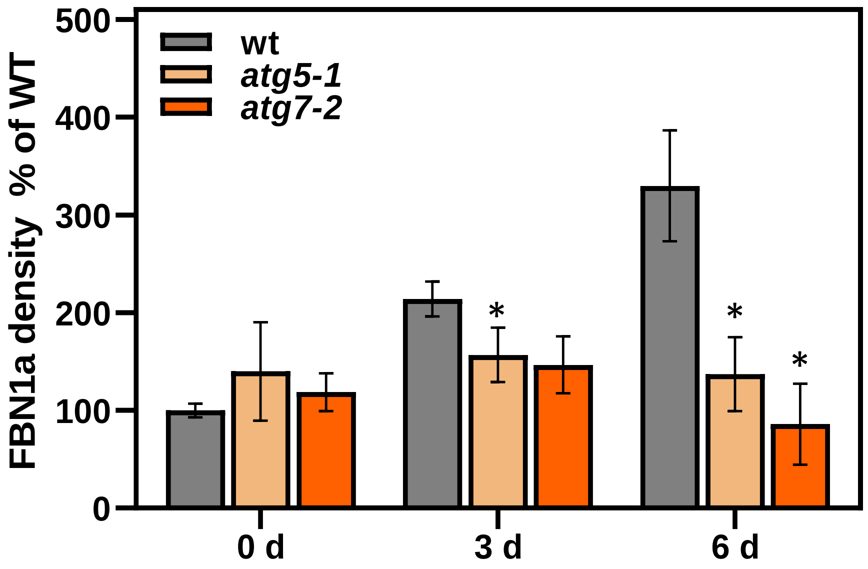


**Supplemental Figure S2: Quantification of the FBN1a protein signal**
Integrated density values for FBN1a were first normalized to loading control and then expressed as a percentage of the obtained value for the wild type (WT) at 0d. Data represent means ±SE; n= 3 biological replicates. An asterisk (*) indicates values that were determined by Student's t-test to be significantly different ($P < 0.05$) from the WT at each time point analyzed.

| | Phospholipids | | | | | |
|---|---|---|---|---|---|---|
| | PC | PE | PG | Galactolipids | Lipid Droplet | β-oxidation |
| *Atatg5* -C | + - | + * | | - | - | + |
| *Atatg5* -N | + | = * | - | - | | + |
| *Zmatg12* -C | - | - | - | - | = | + |
| *Zmatg12* -N | - | - * | - | - | | + |

* Majority of species    Increase    Decrease    Mixed pattern    No change    No data

**Supplemental Figure S3: Compared lipidomics of Arabidopsis and maize *atg* mutants under carbon and nitrogen starvation.** General representation of the changes in Phospholipids, Galactolipds, Lipid Droplet number, and β-oxidation transcripts reported in Arabidopsis and maize under carbon starvation (this study; McLoughlin et al., 2020) and nitrogen starvation (McLoughlin et al., 2018; Have et al., 2019). The scheme does not take into consideration the experimental differences in these studies. Abbreviations- PC: Phosphatidylcholine; PE-: Phosphatidylethanolamine; PG: phosphatidylglycerol; -C: Carbon starvation; -N: Nitrogen starvation

**Supplemental Table S1:** Summary of the main variables of the first three Principal Component (PC) of *atg* mutants and wild type (WT) genotypes after 0 and 6 days of dark treatment. The contribution of which PC is showed as cumulative variance (CV).

| **PC 1** CV = 57.8% | | **PC 2** CV= 69.1% | | **PC 3** CV=78.6% | |
|---|---|---|---|---|---|
| MGDG 34:4 | -0.1813 | TAG 54:9 | -0.2078 | TAG 56:4 | -0.1958 |
| MGDG 34:3 | -0.16 | MGDG 32:3(1) | -0.1916 | TAG 54:6 | -0.184 |
| PC 38:4 (1) | -0.1589 | TAG 52:6 | -0.1681 | TAG 52:4 | -0.1819 |
| MGDG 34:1 | -0.1554 | PC 36:5 | 0.1856 | TAG 54:5 | -0.1765 |

**Supplemental Table S2:** Primers used in the RT-PCR analyses performed in this study

| Gene | Locus | Forward | Reverse |
|---|---|---|---|
| ABC1K7 | AT3G07700 | TCCGTGTTCGGGTTCTTGAGTC | TGGCAACAAGCTGACTTCCTTGG |
| ACT | AT2G37620 | CTTGCACCAAGCAGCATGAA | CCGATCCAGACACTGTACTTCCTT |
| ACX-4 | AT3G51840 | AGGTCAAGCCAGTTTAGGAAAGGC | AATTCCCGACCTAGCGAAGCAG |
| CSY-1 | AT2G42790 | AAGATCATGAGACCACAACAGGTG | TCTCACTGGTGTGTAATGCCTCAG |
| KAT-2 | AT2G33150 | TCCCGTTCTTGGTGTATTCAGGAC | ACCCATGATTGCAGGGTCAACAC |
| MFP-2 | AT3G06860 | ATGCCAATGGGTCCCTTCAGAC | CGATAAACTGCGTTGCGGTTGC |
| PES-1 | AT1G54570 | TGAAGTCTTGGTCCTTGCTAGTGG | AAGTAACCCGTGAAGCCGCTTTG |
| PES-2 | AT3G26840 | AAGGACGCTGGCAACATAAGGG | ACCTGCTGTTTCTATTGGTTTGCC |
| PMDH-1 | AT2G22780 | TGAGCTTCCCTTCTTCGCATCG | TGCCTTCTCTAATCCCATCCTCTC |
| PPH | AT5G13800 | GGTGAGACCGTTATGGGGAAAG | GCATCAGATAGTTCACCACCTCAG |
| SDP-1 | AT5G04040 | TAAGCTCGCGCATCTAGTGGAG | CCGAGCTCTAATACCTGGTTGC |
